# Targeting of the NLRP3 Inflammasome for early COVID-19

**DOI:** 10.1101/2021.02.24.432734

**Authors:** Carlo Marchetti, Kara Mould, Isak W. Tengesdal, William J Janssen, Charles A. Dinarello

## Abstract

Following entry and replication of Severe Acute Respiratory Syndrome-coronavirus-2 (SARS-CoV-2) into ACE2 expressing cells, the infected cells undergo lysis releasing more virus but also cell contents. In the lung, constitutive cytokines such as IL-1α are released together with other cell contents. A cascade of inflammatory cytokines ensues, including chemokines and IL-1β, triggering both local as well as systemic inflammation. This cascade of inflammatory cytokines in patients with COVID-19 is termed “Cytokine Release Syndrome” (CRS), and is associated with poor outcomes and death. Many studies reveal that blocking IL-1 activities in COVID-19 patients reduces disease severity and deaths. Here we report highly significant circulating levels of IL-1β, IL-1 Receptor antagonist, IL-6, TNFα, IL-10 and soluble urokinase plasminogen activator receptor in COVID-19 patients with mild or no symptoms. We also report that in circulating myeloid cells from the same patients, there is increased expression of the NOD-, LRR- and pyrin domain-containing 3 (NLRP3) early in the infection. We observed increased NLRP3 gene expression in myeloid cells correlated with IL-1β gene expression and also with elevated circulating IL-1β levels. We conclude that early in SARS-CoV-2 infection, NLRP3 activation takes place and initiates the CRS. Thus, NLRP3 is a target to reduce the organ damage of inflammatory cytokines of the CRS.

## Introduction

Severe acute respiratory syndrome-coronavirus 2 (SARS-CoV-2) infections manifest as acute lung injury and increased inflammatory response known as coronavirus disease 2019 (COVID-19). Patients with severe symptoms are characterized by unusually high serum inflammatory cytokine levels leading to cytokine release syndrome (CRS) (Del Valle et al. 2020, Lucas et al. 2020) that ultimately results in respiratory distress syndrome and multi-organ failure. CRS contributes to and can be causal in COVID-19 however, the mechanism(s) for initiation of CRS in COVID-19 remains unknown. Numerous trials comparing standard of care in control patients as well as case reports have administered the IL-1 Receptor antagonist (IL-1Ra) anakinra in modest to severe COVID-19 patient, although there are at present no randomized trials. Emerging from these reports is the concept that targeting of IL-1 result in improved outcomes, including deaths. For example, high doses of anakinra reduces deaths as well as number of days in the hospital (Cauchois et al. 2020, Cavalli et al. 2020, Huet et al. 2020). Anakinra has also been administered in less severe hospitalized patients and resulted in similar reduction in disease (Kyriazopoulou et al. 2020). Since anakinra blocks the IL-1 Receptor, the efficacy of anakinra may be due to reducing IL-1α or IL-1β. Other studies report that specifically targeting IL-1β with the neutralizing monoclonal antibody canakinumab also reduces outcomes.

The intracellular processing of IL-1β into its biologically active form is largely governed by cytosolic macromolecular complexes termed inflammasomes (Franchi et al. 2009, Marchetti 2019). Notably, it has been observed that viral proteins of the SARS-CoV virus ORF3a, ORF8b and Viroporin 3a activate the NLRP3 inflammasome (Chen et al. 2019, Shi et al. 2019, Siu et al. 2019). More recently, *in vitro* studies showed that also SARS-CoV-2 induces the NLRP3 inflammasome formation (Xu et al. 2020). Presence of NLRP3 inflammasome aggregates has also been shown in the lungs of fatal COVID-19 pneumonia and in PBMCs and tissues of COVID-19 positive post-mortem patients upon autopsy (Rodrigues et al. 2021, Toldo et al. 2021). Notably, *Rodrigues et al.* have been shown that SARS-CoV-2 virus can infect monocytes leading to the NLRP3 inflammasome formation in these cells (Rodrigues et al. 2021). These studies confirm activation of the NLRP3 inflammasome in COVID-19 in moderate to severe cases. The use of an early treatment for non-hospitalized COVID-19 subjects in order to reduce the burden of hospitalizations and intensive care units is clear (Kim, Read and Fauci 2020). Here we show increased NLRP3 in non-hospitalized SARS-CoV-2 positive subjects. These data support the rationale for early inhibition of NLRP3 to prevent inflammasome formation and the release of IL-1β in SARS-CoV-2 infection.

## Results and Discussion

As shown in Figure 1a, compared to healthy controls, circulating IL-1β is elevated in ambulatory subjects positive for SARS-CoV-2. The mean level in 39 healthy subjects (0.23 pg/mL ±0.03) matches the mean IL-1β (0.33 pg/mL) in 500 healthy Dutch subjects (Ter Horst et al. 2016) whereas in 24 COVID-19 subjects the mean level is 2-fold greater (0.42±0.06; p<0.001). Similarly, mean IL-6 level in infected subjects is greater than 2-fold higher (Figure 1b, p<0.01). The naturally occurring IL-1 Receptor antagonist (IL-1Ra) is 2.5-fold higher in the infected individuals compared to healthy subjects (Figure 1c, p<0.0001). These data are consistent with those of non-infectious autoinflammatory diseases where IL-6 and IL-1Ra are induced by IL-1β and commonly used as surrogate cytokines when circulating levels of IL-1β are exceedingly below most detection assays. Also shown in Figure 1 are significantly elevated levels of tumor necrosis factor alpha (TNFα), IL-10 and urokinase plasminogen activator receptor (uPAR). Plasma uPAR is elevated in COVID-19 patients and levels positively correlate with poor outcomes (Kyriazopoulou et al. 2020).

**Figure 1.**
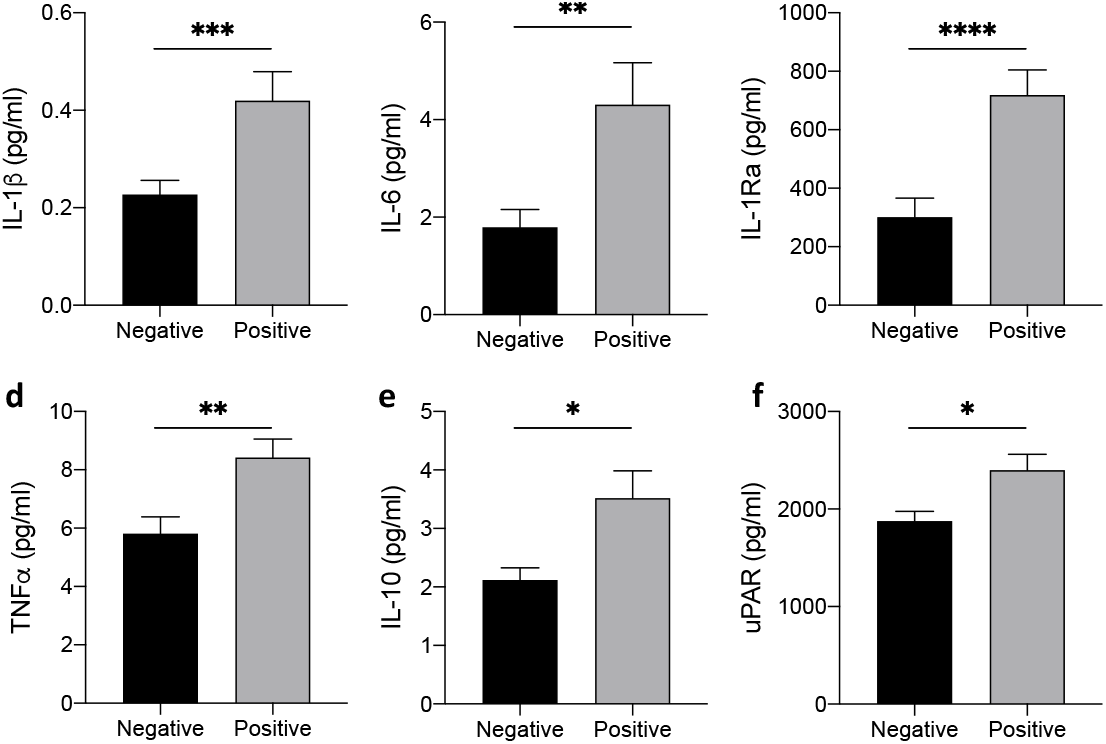
Increased circulating cytokines in early COVID-19. (a-f) Mean ± SEM of plasma IL-1β (a), IL-6 (b), IL-1Ra (c), TNF α (d), IL-10 (e) and uPAR (urokinase plasminogen activator receptor) in SARS-CoV-2 positive patients (N=39) compared to SARS-CoV-2 negative (N=24). *p<0.05; **p<0.01; ***p<0.001; ****p<0.0001.

Since SARS-CoV-2 primes NLRP3 (Rodrigues et al. 2021, Xu et al. 2020), we anticipated an increase in mRNA levels coding for NLRP3 with SARS-CoV-2 infection. We observed a 2-fold increase in NLRP3 levels in buffy coat cells from 27 positive subjects compared to cells from 14 non-infected subjects (Figure 2a). In the same cells, there was a 5.5-fold increase in IL-1β gene expression (Figure 2b). As shown in Figure 2c, NLRP3 and IL-1β mRNA levels are highly correlated (r=0.8, p<0.001) and circulating IL-1β is also positively correlated with NLRP3 gene expression (Figure 2d). Caspase-1 gene expression is elevated 4-fold (Figure 2e), which also correlated with NLRP3 (Figure 2f r=0.8, p<0.001). Thus, in COVID-19 patients the molecular cascade resulting in elevated circulating IL-1β and NLRP3 gene expression is initiated by the infection however, critical for COVID-19 disease is the processing and release of active IL-1β. Using Western blotting, we show evidence of this concept with NLRP3 protein in monocytes from two infected subjects (Figure 2g), but not in cells from an uninfected subject.

**Figure2.**
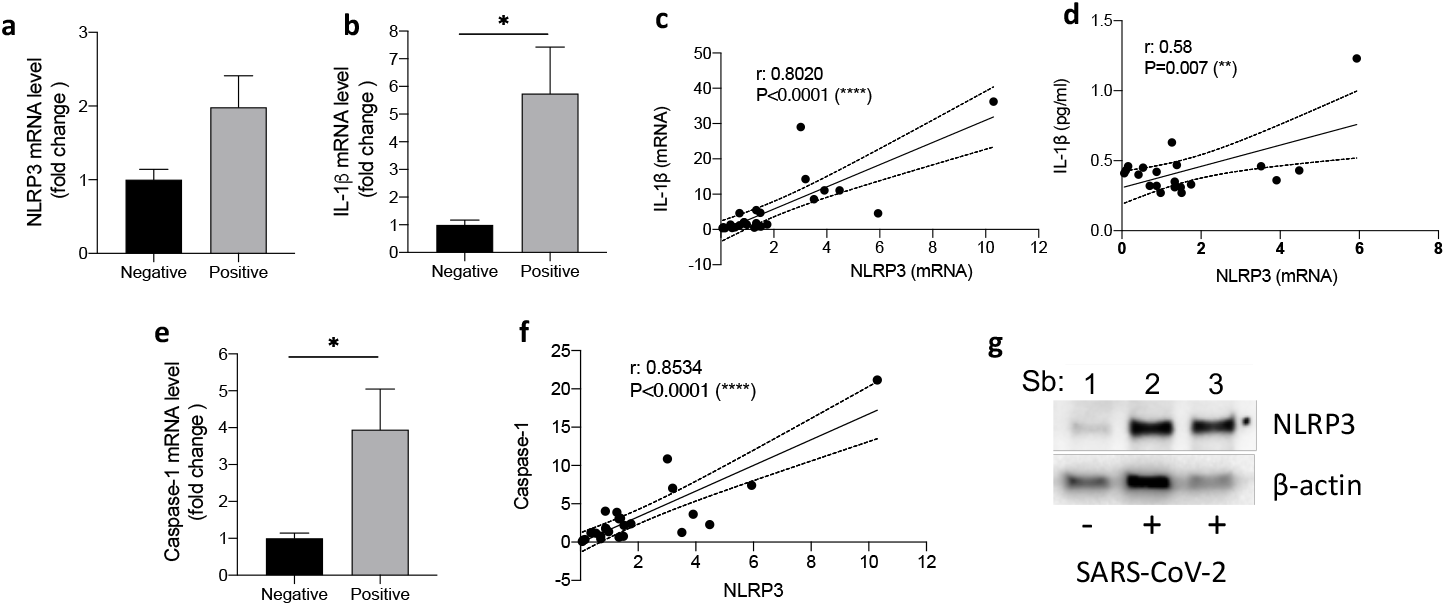
Increased NLRP3 levels in early COVID-19. (a,b) Fold change of NLRP3 (a) and IL-1β (b) mRNA levels from buffy coats of SARS-CoV-2 positive patients (N=27) compared to SARS-CoV-2 negative (N=14) (Wilcoxon signed-rank test). (c) Correlation between NLRP3 and IL-1β gene expression in SARS-CoV-2 positive patients. (d) Correlation between NLRP3 (gene expression) and circulating levels of IL-1β (pg/ml) in SARS-CoV-2 positive patients. (e) Fold change of caspase-1 mRNA levels from buffy coats of SARS-CoV-2 positive patients (N=27) compared to SARS-CoV-2 negative (N=14) (Wilcoxon signed-rank test). (f) Correlation between NLRP3 and caspase-1gene expression in SARS-CoV-2 positive patients. (g) NLP3 protein levels in monocytes isolated from two SARS-CoV-2 positive patients compared to a SARS-CoV-2 negative subject. *p<0.05; **p<0.01; ****p<0.0001.

The morbidity and mortality of COVID-19 often takes place later in the disease when SARS-CoV-2 RNA is absent in secretions as disease worsens and is associated with marked increases in biomarkers of the CRS (Cauchois et al. 2020). Thus, the CRS in COVID-19 is indicative of the organ damaging properties of IL-1β and its downstream cytokines. In the lung, COVID-19 pneumonia is similar to that of Acute Respiratory Disease Syndrome observed in influenza or sepsis. The use of anakinra or canakinumab in COVID-19 disease is associated with recovery and survival. However, it is possible to reduce the detrimental properties of IL-1β and its downstream cytokines by first preventing the processing and release of active IL-1β. The present data provide a rationale to treat patients infected with SARS-CoV-2 early in the course of the disease using a specific NLRP3 inhibitor in order to arrest the progression of IL-1β-mediated CRS. Such a treatment offers an opportunity to reduce hospitalization and the need for supplemental oxygen, particularly in subjects with high risk co-morbidities. At the present time, OLT1177 (rINN dapansutrile) is the only orally active specific NLRP3 inhibitor (Marchetti et al. 2018) that has shown efficacy in two inflammatory diseases, acute gout flares and heart failure (Kluck et al. 2020, Wohlford et al. 2020).

Recent studies have demonstrated that oral administration of colchicine in 4159 non-hospitalized COVID-19 PCR positive patients reduced a composite end-point of hospitalizations and death in 4.6% of treated patients compared to 6.0% of placebo-treated subjects (p<0.04) (Tardif et al. 2021). Colchicine treatment also reduces the risk of cardiovascular events (Nidorf et al. 2020). The mechanism by which colchicine is effective in coronary artery disease as well as in COVID-19 is likely due to reduce IL-1β-mediated inflammation. However, colchicine does not directly inhibit NLRP3 (Hoss and Latz 2018). Unlike specific NLRP3 inhibitors, colchicine affects integrins, cell migration and microtubule assembly.

A significant advantage of targeting the NLRP3 inflammasome is the ability to reduce IL-18 processing. Therefore, specific NLRP3 inhibitors could be used to treat the Macrophage Activation Syndrome (MAS)-like disease in COVID-19, where IL-18 plays a pathological role. Elevated circulating IL-18 correlated with disease severity and poor outcomes in COVID-19 patients (Rodrigues et al. 2021, Lucas et al. 2020). IL-18 is characteristically elevated in non-COVID-19 MAS (Mazodier et al. 2005). Several case series reports that a MAS-like disease develops in COVID-19 patients with markedly elevated levels of D-dimer, which is indicative of MAS in COVID-19 (Aouba et al. 2020). Specific NLRP3 inhibition will reduce both IL-1β as well as IL-18 and thus targets two agonists of COVID-19 disease.

## Methods

### PBMCs

Peripheral blood mononuclear cells (PBMCs) were isolated from drawn blood by gradient centrifugation using Ficoll-Paque (Pharmacia Biotech, Uppsala, Sweden). PBMCs were suspended in Roswell Park Memorial Institute 1640 medium supplemented with 50 µg/mL gentamicin, 2 mM glutamine, and 1 mM pyruvate and cultured for 24 hours.

### Cytokines measurements

Plasma levels of IL-1β, IL-6, IL-10 and TNFα were measured with the Ella platform (Protein Simple, San Jose, CA, USA) using multiplex cartridges. Soluble uPAR was determined using Quantikine ELISA (R&D Systems, Minneapolis, MN, USA).

### Gene expression

RNA was isolated according to the manufacturer’s protocol (Thermo Fisher Scientific) and synthesized into cDNA using SuperScript III First-Strand (Thermo Fisher Scientific). Quantitative PCR (qPCR) was performed on cDNA using Power SYBR Green PCR master mix (Thermo Fisher Scientific) on Bio-Rad (Bio-Rad Laboratories, Hercules, CA) CFX96 Real time system. Gene expression was carried for the following mRNAs: *nlrp3, caspase1* and *il1b* with *gapdh* used as reference gene using the following primers:

*nlrp3*: GAATCTCAGGCACCTTTACC and GCAGTTGTCTAATTCCAACACC
*caspase1*: AAGTCGGCAGAGATTTATCCA and GATGTCAACCTCAGCTCCAG
*il1b*: CTAAACAGATGAAGTGCTCCTTCC and CACATAAGCCTCGTTATCCCA

### Western blotting

Cells were lysed using RIPA buffer supplemented with protease inhibitors (Roche), centrifuged at 13,000 *g* for 20 min at 4°C and the supernatants were obtained. Protein concentration was determined in the clarified supernatant using Bio-Rad protein assay (Bio-Rad Laboratories, Hercules, CA). Proteins were electrophoresed on Mini-PROTEAN TGX 4−20% gels (Bio-Rad) and transferred to nitrocellulose 0.2μM (GE Water & Process Technologies). Membranes were blocked in 5% dried milk in PBS-T 0.5% for 1 hour at room temperature. Primary antibodies for NLRP3 1:1000 (Adipogen) was used in combination with peroxidase-conjugated secondary antibodies and chemiluminescence to detect the protein concentration. A primary antibody against β-actin (Santa Cruz Biotechnology) was used to assess protein loading.

### Statistical analysis

Significance of differences was evaluated with Student’s two-tail T-test using GraphPad Prism (GraphPad Software Inc, La Jolla, CA) or Wilcoxon signed-rank test as indicated. For the correlation studies the distribution were computed using Pearson correlation coefficients and Statistical significance was calculated with two-tailed option with the confident interval set at 95%. Statistical significance was set at p<0.05.

